# High-efficiency PET Degradation with a Duo-enzyme System Immobilized on Magnetic Nanoparticles

**DOI:** 10.1101/2024.04.15.589601

**Authors:** Siddhi Kotnis, Siddhant Gulati, Qing Sun

## Abstract

The widespread consumption of PET worldwide has necessitated the search for environment-friendly methods for PET degradation and recycling. Among these methods, biodegradation stands out as a promising approach for recycling PET. The discovery of duo enzyme system PETase and MHETase in 2016, along with their engineered variants, has demonstrated significant potential in breaking down PET. Previous studies have also demonstrated that the activity of the enzyme PETase increases when it is immobilized on nanoparticles. To achieve highly efficient and complete PET depolymerization, we immobilized both FAST PETase and MHETase at a specific ratio on magnetic nanoparticles. This immobilization resulted in a 2.5-fold increase in product release compared with free enzymes. Additionally, we achieved reusability and enhanced stability of the enzyme bioconjugates.

## 1. Introduction

Polyethylene terephthalate (PET) is a widely used thermoplastic for packaging, constituting up to 67% of the total packaging plastic in the US.^1^ Due to its resistance to chemical and mechanical changes, it is a significant challenge to manage PET waste efficiently.^2^ The two major ways of recycling and upscaling PET are through chemical and mechanical recycling.^3^ Chemical recycling is achieved through solvolysis or pyrolysis where PET is degraded to its monomers of ethylene glycol (EG) and terephthalic acid (TPA).^4^ These processes are economically unfeasible and involve harsh reaction conditions, leading to the release of harmful toxins into the environment.^5,6^ On the other hand, mechanical recycling reshapes PET into other desirable shapes, but this can only be done for a limited number of cycles before the polymer loses mechanical strength and cannot be used further.^7^

An environmentally friendly and cost-effective alternative is the biodegradation of PET using enzymes as biological catalysts to degrade PET into its monomers.^8,9^ Among these enzymes, PETase exhibits high activity at room temperature, making it ideal for PET degradation at mild reaction conditions.^10^ Following the discovery of PETase in 2016, there have been many mutations of PETase showing beber activities and thermostabilities,^11–13^ within which FAST PETase exhibited the best performance with 10 times beber activity than wildtype PETase at 30°C. Moreover, PETase, along with MHETase, shows a synergistic effect in degradation of PET to monomers TPA and EG.^14^ The presence of MHETase, even small ammount, is crucial for efficient complete depolymerization of PET.^14^

One of the major obstacles to commercialize the enzyme system is the enzyme aggregation and instability at higher concentration.^15^ Reports indicate that at concentrations exceeding 150nM, PETase begins to aggregate, resulting in reduced enzymatic efficiency.^16^ To overcome these challenges, enzyme immobilization has emerged as an attractive alternative to solve the enzyme aggregation and instability problem.^17–20^ Magnetic nanoparticles, within all different supporting plaeorms for enzyme immobilization^17^, have advantages such as being biofriendly and easy separation.^21–23^ Immobilization of single enzyme PETase on nanoparticles including cobalt phosphate nanoflowers and iron oxide nanoparticles has shown increased PET degradation efficiency.^24,25^ However, these system has not achieved complete PET degradation due to the limitations of the single-enzyme approach.

In this work, we propose a novel approach of immobilizing both FAST PETase and MHETase on iron oxide nanoparticles in appropriate ratio to maximize PET degradation and achieve complete depolymerization of PET to TPA and EG. These bioconjugates, formed through the electrostatic interactions between his tag on the enzymes and the nanoparticle,^22^ enhances the efficiency of the PET degradation system. Additionally, the bioconjugates exhibit improved stability and reusability, demonstrating the potential of the immobilization system for plastic waste management.

## 2. Experimental section

### 2.1 Expression and Purification of proteins

pBbE8k-FAST PETase was constructed and transformed on *E. coli* NEB5α and then on *E. coli* ShuffleT7 for expression. pET21a-MHETase was constructed and transformed on *E. coli* NEB5a and then *E. coli* C41(DE3) for expression. The enzymes were expressed and then purified using the hexa-histidine tag with Ni-NTA column.

### 2.2 Immobilize MHETase and FAST PETase on MNPs

Varying concentrations of the enzyme mixture(0.1-0.3g/L) were added to 1g/L Magnetic Nanoparticles (Iron Oxide Nanoparticles, 15nm, Sigma Aldrich) and incubated at 25°C with shaking at 250 RPM for 2h. There are two approaches to addition of enzymes on the MNPs, simultaneous and sequential. The simultaneous approach involves addition of both enzymes at the same time, while the sequential approach involved addition of MHETase at t=0h and then addition of FAST PETase at t=1h. The MNP-enzyme nano-bioconjugates are separated using a Magnet. The supernatant was analyzed for quantifying the protein through DC Assay (Biorad). A 10% SDS PAGE Gel is also run with the supernatant to visualize the enzyme immobilized on the MNPs. The nano-bioconjugates were washed thrice with PBS and stored in PBS at 4°C.

### 2.3 Enzyme activity on PET film

PET film (Goodfellow ES301445) was washed with 70% Ethanol. 1uM total enzyme was incubated with the PET film to a final volume of 300uL reaction medium (pH 7.5 50 mM NaH2PO4, 100 mM NaCl, 10% (v/v) DMSO) for 96h at 30°C and 250 RPM. Samples were collected at 96h, and heat treated at 85°C for 20 minutes to denature and inactivate enzymes in the solution. The samples were passed through 0.22um spin filters before HPLC analysis. The TPA concentration was then determined through HPLC analysis at 240nm. Same steps are followed for bioconjugates as well, where NBCs are added to achieve a final enzyme concentration of 1uM.

### 2.4 Reusability and Stability

The MNPs were separated from the reaction medium using a magnet. They were washed thrice with PBS (pH 7.4) and stored at 4°C. *p*NPA (*para*-Nitrophenol acetate) was used as a substrate for the determination of activity of the bioconjugates and free enzyme for reusability and stability. For the assay, a final concentration of 2mM *p*NPA and 5% v/v DMSO was used with a 200uL final reaction volume of PBS (pH 7.4).^26^ The assay is conducted in a 96 well plate and the absorbance is monitored at 405 nm for 10 minutes with 30 second intervals.

## 3. Results and discussion

### 3.1 Immobilization of enzymes on MNPs

PETase and MHETase have shown a synergistic effect in the complete depolymerization of PET to its monomers of TPA and EG.^14^ The presence of MHETase, even small ammount, is crucial for efficient complete depolymerization of PET. However, the concentration of MHETase has mininal impact on the depolymerization performance of the two enzyme system.^14^ With the mutant varient FAST PETase, we observed a similar behavior, achieving optimal product yield at a molar ratio of 20:1 between FAST PETase and MHETase. At MHETase concentrations lower than this, the product release began to decline (Supplementary Fig. 1).

Aiming to maximize PET degradation, we sought to immobilize the enzymes on nanoparticles at the optimal ratio of 20:1. Given the relatively minor ratio of MHETase, we employed two immobization strategies to maximize enzyme loading at this ratio: a simultaneous method, where both enzymes were added together, and a sequential method, where MHETase was added before FAST PETase.

The immobilization of FAST PETase and MHETase onto nanoparticles was verified qualitatively using SDS PAGE analysis (Supplementary Fig. 2) and quantitatively using protein centration DC Assay. The analysis of the supernatant before and aper magnetic beads immobilization indicated the successful enzyme binding onto the nanoparticles Figure (1b). We explored enzyme concentrations ranging from 0.1g/L to 0.3g/L for both immobilization methods. While sequential addition showed no advantages at the lower concentration of 0.1g/L, it displayed increased enzyme loading at 0.3g/L, likely due to reduced competitive binding and steric hindrance of enzymes as compared to simultaneous addition. At an initial concentration of 0.3 g/L, enzyme loading for sequential addition (∼0.2g/L) was approximately double that of simultaneous addition (∼0.1g/L). Notably, sequential addition resulted in a higher proportion of MHETase on the nanoparticles, as evidenced by SDS-PAGE analysis of the supernatants (Supplementary Fig. 2). Since highest enzyme loading was observed for the initial total enzyme concentration of 0.25g/L and sequential addition was favored for higher percentage of MHETase immobilization, we used these conditions for the subsequent activity assays.

### 3.2 Activity on PET film

Bioconjugates, formed using sequential addition with an enzyme concentration of 0.25g/L, were tested for enzyme activity on PET films by High-Performance Liquid Chromatography (HPLC) analysis of depolymerization products. TPA were analyzed using HPLC traces at a wavelength of 240nm (UV) (Supplementary Fig. 3). The peak areas were quantified against a TPA calibration curve (Supplementary Fig. 4). Comparing bioconjugates to an equivalent free enzyme system, the bioconjugates released approximately 8mM of TPA, significantly higher than the 3mM observed from the free enzyme system (Fig. 2). This enhanced activity is likely due to the increased stability of enzymes, which prevents aggregation above a concentration of 150nM, as well as the synergy between FAST PETase and MHETase from close proximity, which has shown to be beneficial. ^14^

**Figure 1:**
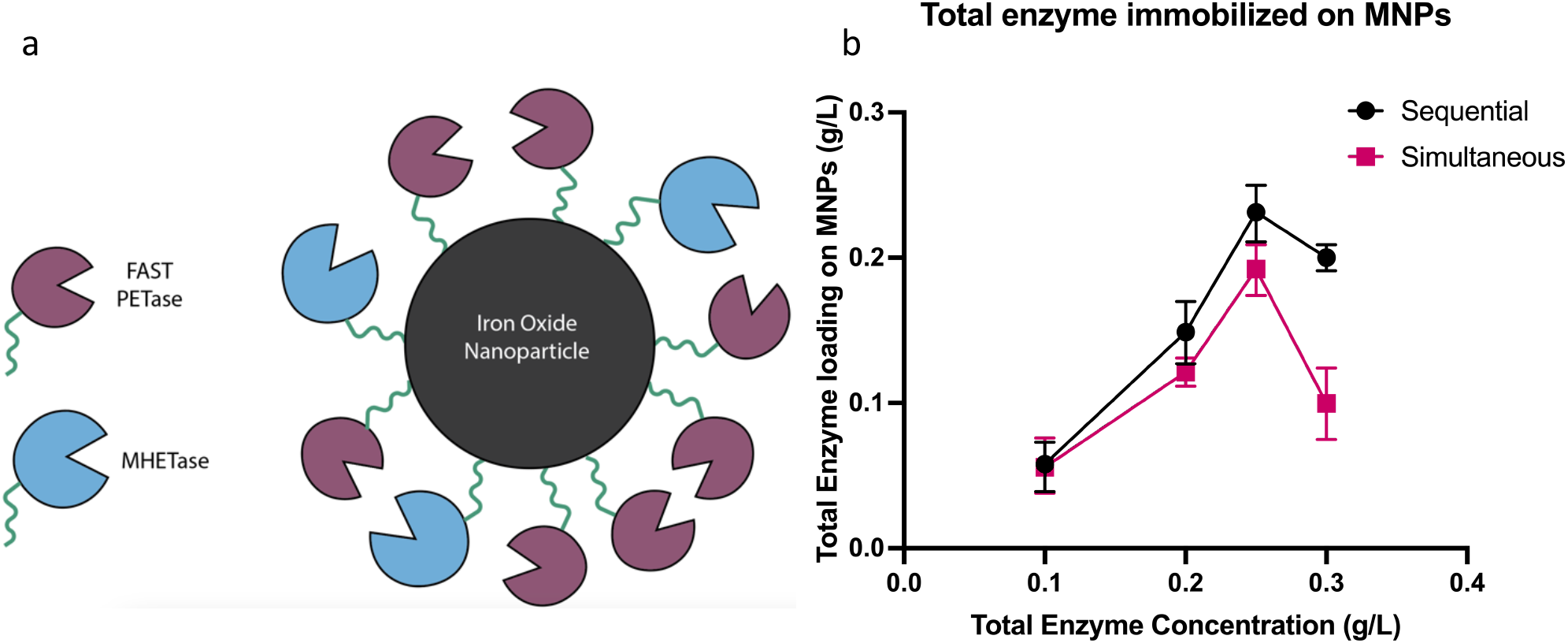
Overall scheme of the paper and total enzymes immobilization on magnetic nanoparticles. (a). FAST PETase and MHETase immobilizing on the Iron oxide nanoparticle through a histag. (b). The comparison in enzyme loading for sequential (black) and simultaneous (pink) addition of enzymes.

**Figure 2.**
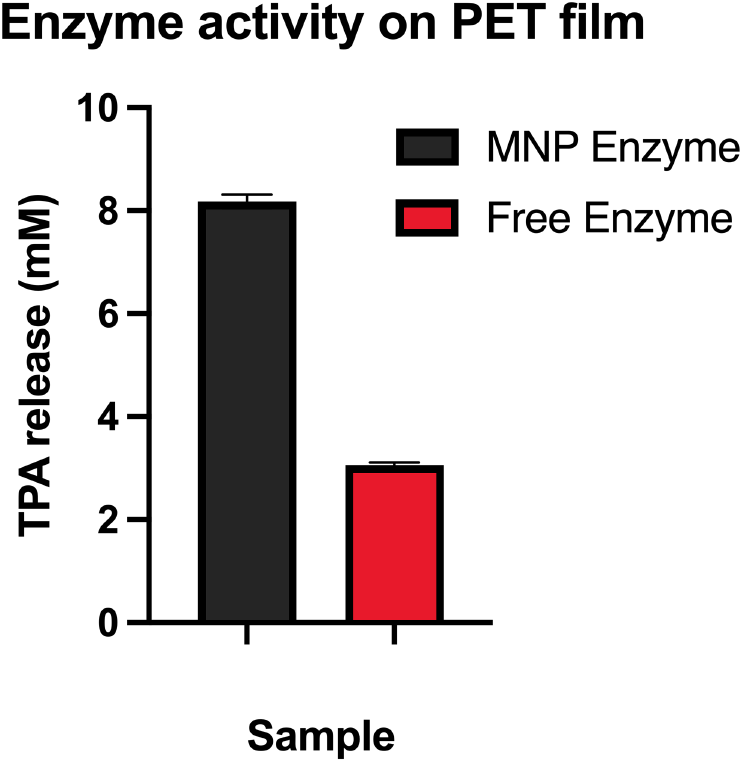
Comparison of TPA released for bioconjugates and free enzyme samples.

Notably, since the bioconjugates have both FAST PETase and MHETase, we observed complete depolymerization of PET to TPA without detecting reaction intermediates like BHET or MHET in the chromatogram of HPLC traces (Supplementary Fig. 3). Furthermore, we observed a product release of 8mM TPA after a 96h incubation of PET film at 30°C which is a much higher degradation rate than what has been reported so far. ^24–26^

### 3.3 Recyclability

We further tested the recyclability of the bioconjugates through magnetic separation. Instead of PET depolymerization, we used *para*-nitrophenyl acetate (*p*NPA), an alternative substrate for esterase, which upon hydrolysis, yields *para*-nitrophenol – a compound detectable by its yellowish-green color and quantifiable at 405nm absorbance. Through successive cycles, we found that the bioconjugates retain over 80% of their initial activity for the first four cycles, with a subsequent decline to 50% activity aper cycle 4, remaining active up to cycle 10 (Fig. 3a). Control samples washed with PBS (pH 7.4) showed the same decline trend in activity, suggesting that the activity loss was not due to enzyme deactivation or substrate inhibition, but potentially from agglomeration and deagglomeration of the bioconjugates during magnetic separation (Fig. 3a). ^24^ Despite some loss of activity, the bioconjugates demonstrated significant reusability for up to ten cycles, with minimal effort required for magnetic separation.

**Figure 3.**
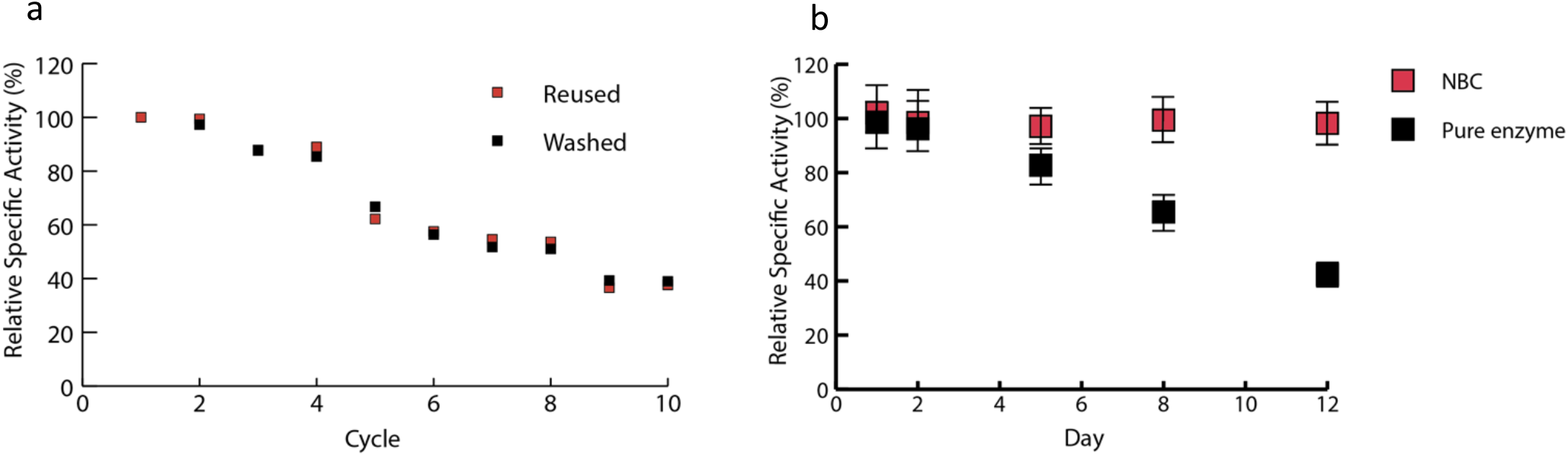
Reusability and stability of bioconjugates. (a). Reusability of enzymes for reused samples and control washed samples; (b) Stability of bioconjugates and free enzyme samples.

### 3.4 Stability of bioconjugates

The stability of bioconjugates was assessed by comparing the activity of enzymes between bioconjugates and equivalent free enzymes, both stored at 4°C. To measure enzyme activity post-storage, *p*NPA assay was used. The bioconjugates containing both FAST PETase and MHETase showed stability for more than a week without any significant loss (Fig. 3b). Conversely, the free enzymes, stored at the same temperature, showed a gradual decrease to approximately 40% of their initial activity aper a 12-day period.

## 4. Conclusion

The immobilization of enzymes onto inorganic scaffolds has proven effective to enhance enzyme activity. In our study focusing on degrading PET using dual-enzyme system, we opted to immobilize FAST PETase and MHETase onto iron oxide nanoparticles. These two enzymes work in synergy to completely depolymerize PET to its monomers. Through sequential addition, we successfully immobilized these enzymes on the iron oxide nanoparticles, ensuring an excess of FAST PETase enzyme. This sequential addition approach not only facilitated efficient immobilization but also improved the total enzyme loading on the nanoparticles compared with simultaneous addition. This two-enzyme system on nanoparticles exhibited a 2.5-fold increase in TPA formation compared with free enzyme system. Furthermore, we demonstrated the reusability of the enzyme bioconjugates using simple magnetic separation. Additionally, the enzyme bioconjugates showed enhanced stability when stored at 4°C compared to free enzyme. Overall, we achieved complete depolymerization of PET using our enzyme bioconjugates, along with additional advantages of reusability and stability through immobilization on nanoparticles. This work could apply to other multi-enzyme systems with non-equimolar requirement to enhance activity through immobilization.

## Credit Author Statement

Q.S. did the conceptualization. S. K. designed and conducted the experiments. S. G. helped with earlier constructs and experiment assay. S. K. drafted the manuscript. Q. S. revised the manuscript and did the funding acquisition.

## Acknowledgements

This work was supported by funds provided by a National Science Foundation (NSF) grant to Q. S. (2203715), USA; NSF Emerging Frontiers in Research and Innovation (EFRI) to Q. S. (2132156), USA; Texas A&M Excellence Fund X-grants to Q. S., Texas A&M, USA; Texas A&M Targeted Proposal Teams (TPT) grant to Q. S., Texas A&M, USA.

## Supplementary Figures

**Figure 1.**
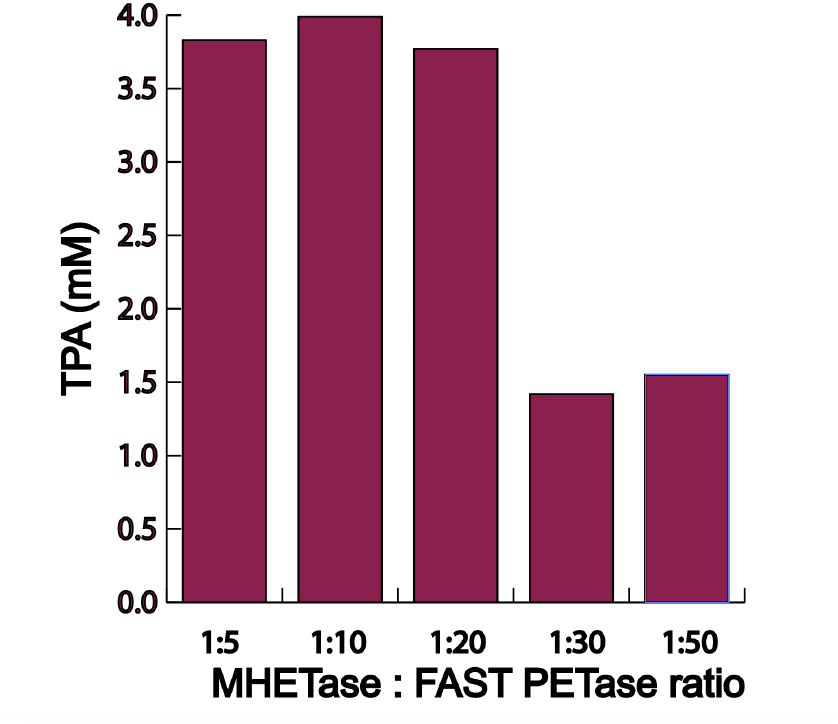
TPA product release for varied MHETase : FAST PETase ratios

**Figure 2.**
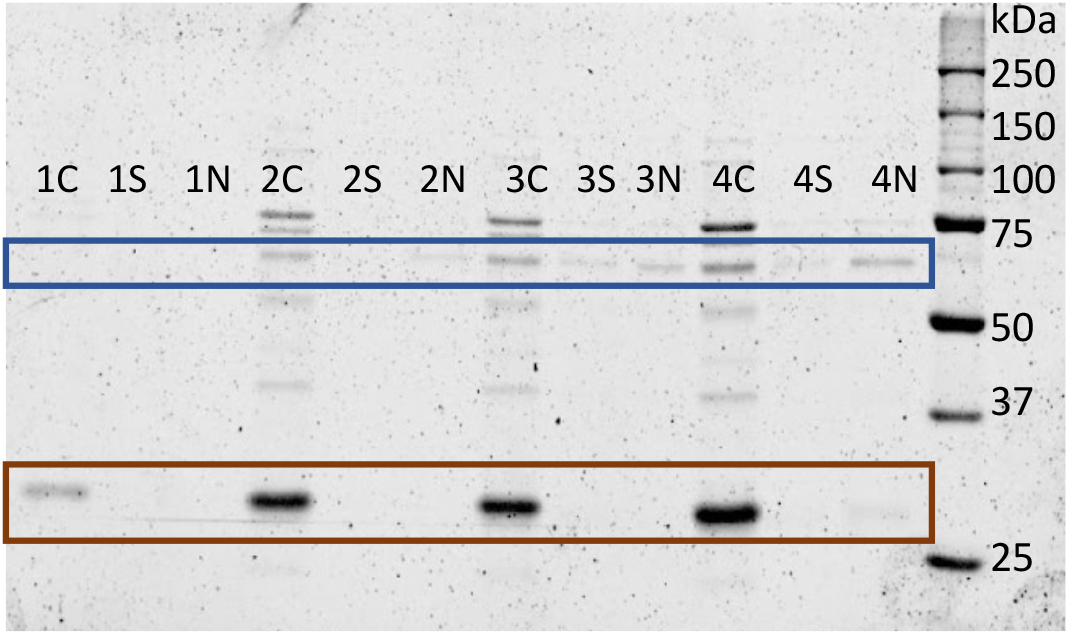
SDS PAGE Gel for comparing sequential and simultaneous addition of enzyme. (1-0.1g/L, 2-0.2g/L, 3-0.25g/L, 4-3g/L. C represents control, or total enzyme at the time of addition. S represents supernatant from sequential addition. N represents supernatant from simultaneous addition)

**Figure 3.**
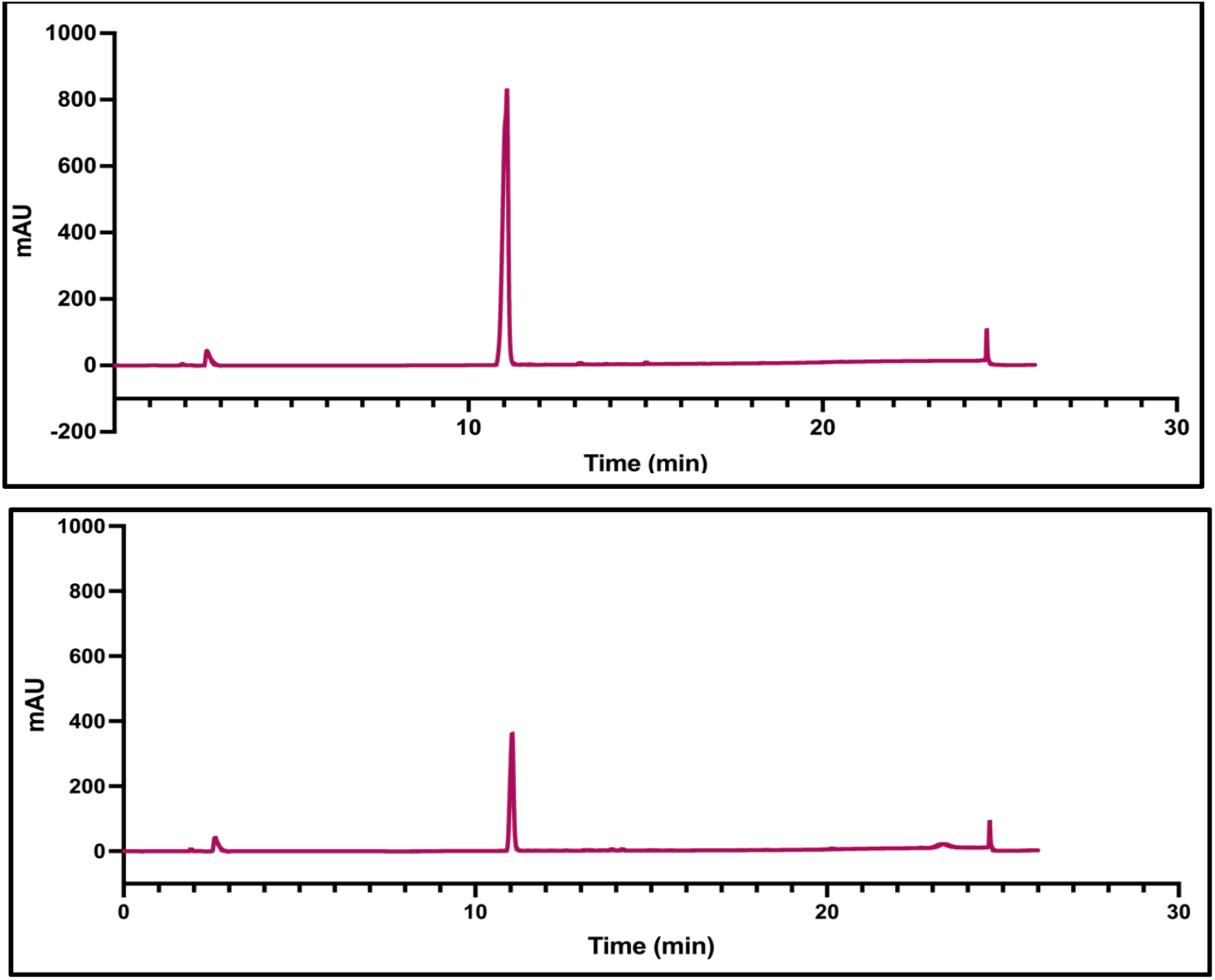
Chromatograms showing peaks of degraded product TPA

**Figure 4.**
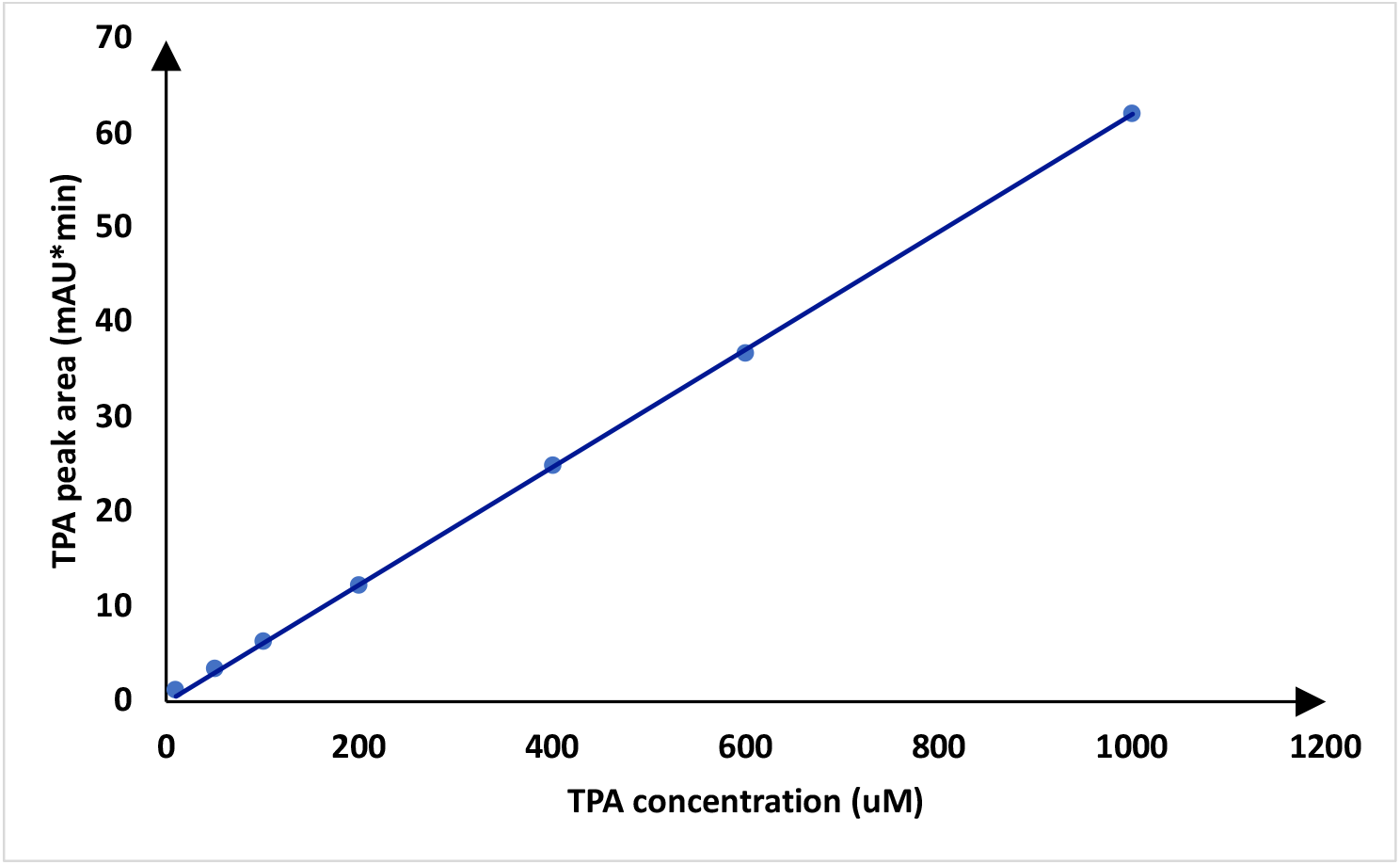
TPA Calibration curve

